# HetNetEX: Exact Asymptotic Inference in Heterogeneous Biomedical Knowledge Graphs

**DOI:** 10.64898/2026.07.05.736581

**Authors:** Tusharkanti Ghosh, Lucas A. Gillenwater, Casey S. Greene, James C. Costello

## Abstract

Heterogeneous biomedical knowledge networks (hetnets) integrate disparate data types, drugs, genes, diseases, and pathways, across independent sources; Hetionet (https://het.io) is a widely used example. A standard approach for assessing connectivity significance is XSwap, which permutes the hetnet *P* times and fits a gamma-hurdle null model to the degree-weighted path count (DWPC), pooling permuted values across pairs with matching source and target degrees to increase the effective sample size. This permutation approach has been highly successful in practice, but it faces four practical constraints in large graphs: (1) a finite resolution for the smallest reportable *p*-values, (2) computational cost that grows prohibitive at path lengths *L* ≥ 4 or 5, (3) a variance model (Var ∝ *µ*^2^) that departs from the configuration-model form (1 + *κ*)*µ*, and (4) *O*(*P* 10*m L*) runtime. To complement this approach, we present HetNetEX (Heterogeneous Network EXact inference), which computes the null DWPC distribution analytically from degree sequences using the configuration model in *O*(*Ln*) time. In simulations at *P* = 200 across *L* = 1–4, HetNetEX achieves Spearman *ρ >* 0.96 concordance with XSwap rankings while being *>*10,000× faster and providing analytical *p*-values without a resolution ceiling. High-degree pairs show larger XSwap sampling error than low-degree pairs, reflecting the finite-sample nature of permutation that analytical computation avoids.

## 1 Introduction

Heterogeneous biomedical knowledge networks (hetnets) are graphs whose nodes represent diverse features, compounds, genes, diseases, pathways, and whose edges represent typed, often directed and weighted, relationships. Unlike homogeneous graphs, hetnets encode the relational structure of biology (a compound *binds* a gene, a gene *participates in* a pathway). This typed structure enables metapath-based reasoning: traversing specific sequences of edge types rather than performing exhaustive path search, which shrinks the search space and filters to relations relevant to the biological question (e.g., Compound → Gene → Pathway rather than all 3-hop paths) [2].

Hetionet v1.0 contains 47,031 nodes spanning 11 types (1,552 compounds, 20,945 genes, 137 diseases, 1,822 pathways) and 2,250,197 edges spanning 24 types from 29 databases [1]. The Rephetio project used Hetionet to predict drug repurposing opportunities, achieving strong cross-validated performance (AUROC ≈ 0.97) in recovering known drug indications [1]. Central to this is the degree-weighted path count (DWPC), which we now define.

### 1.1 Degree-Weighted Path Count

The DWPC quantifies connection strength between a source *s* and target *t* along a metapath (*e*_1_, …, *e*_*L*_):

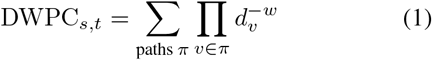

where *d*_*v*_ is the degree of node *v* and *w* = 0.4 down-weights paths through high-degree hubs [2]. The rationale reflects how biomedical data are generated: high-degree nodes accumulate connections through a mixture of systematic assays and literature-based curation, and these non-random, non-biological processes introduce degree-driven biases unrelated to true signal [2]. TP53 (degree *>* 500) accrues paths largely because it is heavily studied, whereas COX2 (degree 12) contributes paths more likely to reflect specific relationships; DWPC weighting discounts paths through high-degree intermediates. For example, from Aspirin (*d*=4) to Inflammation (*d*=8): the path through COX2 (*d*=12) contributes 4^*−*0.4^ · 12^*−*0.4^ · 8^*−*0.4^ = 0.091, while the path through TP53 (*d*=500) contributes only 0.019; the hub path is discounted 5×, matching the known pharmacology of aspirin on COX2 [3].

### 1.2 The Significance Problem

An observed DWPC of 0.110 is uninterpretable without a reference: it could be significant if the degree structure predicts near-zero connectivity, or unremarkable if *s* and *t* are both hubs where many paths arise from the same curation- and assay-driven biases rather than genuine biology. A null model provides this reference by computing the DWPC distribution under degree-preserving randomization (graphs that retain each node’s degree but destroy specific wiring), and a *p*-value quantifies how unlikely the observed value is.

The XSwap algorithm [6] generates degree-preserving random graphs via Markov-chain edge swapping, applied separately per edge type for hetnets [2]. Whereas Rephetio used only *P* = 5 permutations for aggregate features, the connectivity search application [4] targets null distributions for *individual* DWPCs, which needs far more permuted values. Himmelstein et al. therefore generated *P* = 200 permuted hetnets and further augmented the effective sample size by grouping DWPC values according to source and target degree: because permutation preserves only node degree, values among pairs with the same (*d*_*s*_, *d*_*t*_) are exchangeable and serve as additional pseudo-permutations. XSwap then fits a gamma-hurdle model to this pooled null, separating the zero component from a gamma-distributed positive component.

Zietz et al. [5] introduced the XSwap Edge Prior, which analytically computes single-edge existence probabilities. It is restricted to *L* = 1: at *L* ≥ 2, edge independence is violated because paths through a shared hub *v* are positively correlated through *v*’s degree constraint, a dependence captured by our overdispersion *κ* (Theorem 2). It also assumes the XSwap chain has mixed; for dense edge types (GiG, *m* = 147,164) the high rejection rate (∼ 80%) can slow mixing.

### 1.3 Constraints of Permutation-Based Inference

The permutation approach has been remarkably effective: it is nonparametric in its sampling stage, targets exactly the null of interest, and, through degree-grouping, extracts far more resolution than a naive 1*/*(*P* +1) bound would suggest. The points below are not failures of XSwap but intrinsic properties of any finite-sample permutation scheme that an analytical method can complement.

#### Finite *p*-value resolution

A single pair’s empirical *p*-value is bounded below by 1*/*(*P* +1) = 0.005 at *P* = 200. Degree-grouping [4] substantially improves this: pooling exchangeable pairs with the same (*d*_*s*_, *d*_*t*_) makes the effective sample size, and hence resolution, many times larger than *P* for common degree combinations. Resolution is thus not a fixed floor but varies with how many pairs share a degree combination; for the rarest combinations (including some high-degree hub pairs of interest), fewer pairs are available and resolution is coarser. Analytical evaluation removes this dependence on sample size.

#### Computational cost at long paths

XSwap runs in *O*(*P* .10*m* .*L*). On Hetionet, *L* = 5 takes ∼8 hours per metapath and *L* = 8 extends to years (Table 1), placing biologically meaningful long paths out of reach.

**Table 1.** Computational complexity: XSwap vs HetNetEX.

|  | XSwap | HetNetEX |
| --- | --- | --- |
| Complexity | $O(P \times 10m \times L)$ | $O(L \times n)$ |
| **Hetionet GiG** ($m = 147,164$, $n = 20,945$, $P = 200$): | | |
| Swap attempts | **294M** | 0 |
| Rejected | $\sim 235\text{M}$ ($\sim 80\%$) | 0 |
| $L = 1$ | $\sim 820$ s | **0.02 s** |
| $L = 4$ | $\sim 30,000$ s | **0.05 s** |
| $L = 8$ | $\sim 3.4$ yr | **0.08 s** |
| **Speedup** | $\sim 40,000 \times (L=1) \rightarrow \mathbf{1.3B} \times (L=8)$ | |

$m \gg n$ in biological graphs, so XSwap (scaling with $m$ ) is far costlier than HetNetEX (scaling with $n$ ). Platform: Apple M2 Pro, 64 GB RAM.

#### Variance model mismatch

The gamma-hurdle implies Var ∝ *µ*^2^*/k* (quadratic), whereas the configuration-model variance is (1 + *κ*)*µ* (linear; Theorem 2). Where these diverge, the fitted null misstates the spread, overstating it for high-*µ* hubs (conservative) and understating it for low-*µ* pairs (anti-conservative).

#### Computational overhead

At Hetionet scale, XSwap’s chain performs ∼294M swap attempts per query; ∼ 80% are rejected to preserve degrees, a necessary part of mixing rather than wasted work. HetNetEX avoids this by computing directly from degree sequences, with cache-friendly sequential dot products.

### 1.4 Contributions

We present **HetNetEX** (**Het**erogeneous **Net**work **EX**act inference), which computes the null DWPC distribution analytically. It targets the same null as XSwap; where XSwap samples with *P* iterations, HetNetEX computes from closed-form degree formulas, and as *P* →∞ XS wap converges to HetNetEX (Theorem 5). Contributions: (i) closed-form *µ* and overdispersion *κ* in *O*(*Ln*) (Theorems 1–2); (ii) Poisson (*L* ≤ 3) and CLT (*L* ≥ 4) distributional theory (Theorems 3–4); (iii) proof of equivalence to the XSwap stationary distribution (Theorem 5); and (iv) empirical validation at *P* = 200, *L* = 1–4, characterizing where finite-sample permutation and analytical inference diverge.

## 2 Methods

### 2.1 Problem Formulation

Let *G* = (*V, E*) have edge types T_*E*_; for *e* ∈ T_*E*_, *m*_*e*_ is the edge count and 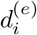 the degree of node *i*. A metapath P = (*e*_1_, …, *e*_*L*_) is a sequence of *L* edge types. Given DWPC_obs_(*s, t*; *P*), we compute *p* = Pr(*Y* ≥ DWPC_obs_| null), where *Y* is the DWPC in a degree-preserving random graph. XSwap estimates *p* by generating *P* graphs and counting exceedances; HetNetEX derives the null mean *µ* and spread *σ* (via *κ*) from degree sequences and evaluates the tail analytically, Poisson CDF when *µ <* 10, Normal when *µ* ≥ 10.

### 2.2 Assumptions

#### Assumption 1

(Configuration model). *For each edge type e, the null graph is drawn from G* (**d**^(*e*)^): *uniform half-edge matching preserving degrees [7, 8]. This is the same null XSwap targets*.

#### Assumption 2

(Edge-type independence). *Edge types are independent given node identities, since they come from separate databases (DrugBank [9], STRING [10], Reactome [11]). Whether a compound binds a gene does not affect whether that gene participates in a pathway. Conditioning on node identities accounts for shared nodes; the assumption concerns only whether* wiring *within one edge type affects another*.

#### Assumption 3 (Sparsity)

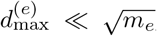, *giving* 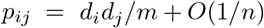 *This is not strictly satisfied for extreme hubs (TP53: d*_max_ = 542, 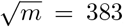, *ratio 1*.*42), but TP53 is a singular outlier: the 99th-percentile degree is* ∼ *60 (ratio 0*.*16) and the median is 7 (ratio 0*.*018). The per-pair error is* 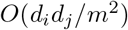, *<* 0.2% *even for TP53*.

### 2.3 Theoretical Results

#### Theorem 1

(Expected null DWPC). *Under Assumptions 1–3*,

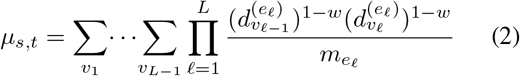

*computable in O*(*Ln*) *via a rank-1 matrix chain* 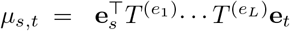

*Proof*. With *I*_*π*_ the path-existence indicator and 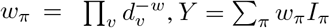 and E[*Y*] = ∑ _*π*_ *w*_*π*_E[*I*_*π*_]. Under the configuration model, edges on distinct pairs are asymptotically independent [8], so 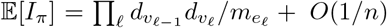. Combining *d*^*−w*^ with the configuration factor *d* yields *d*^1*−w*^. Each 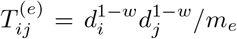 has rank 1, so the chain is *L* dot products in *O*(*Ln*) rather than *O*(*Ln*^2^).

#### Corollary 1

*w enters only through* 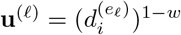; *the framework holds for any w* ∈ [0, 1].

#### Theorem 2

(Variance and overdispersion). Var(*Y*) ≈ (1 + *κ*)*µ where* 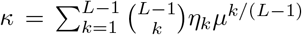 *and is the degree heterogeneity ratio*.

*Proof*. With *λ*_*π*_ = *w*_*π*_E[*I*_*π*_], Var(*Y*) = ∑_*π*_ *λ*_*π*_(1 − *′λ′π*) + ∑ _*p* ≠ *π*_Cov(*I′π, I′π*). Paths sharing no intermedi are ways to share *k* positions, each weighted by the ate node are independent; paths sharing *k* of *L* − 1 positions contribute covariance ∝ *n*^*−*(*L−k*)^*λ*_*π*_*λ*_*π*_*′*, and there are 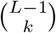 ways to share *k* positions,each weighted by the second moment *η*_*k*_. When *κ* = 0 (no shared intermediates, or homogeneous degrees *η*_*k*_ = 1), Var = *µ* (Poisson). Real networks are skewed (*η*_*k*_ ≫ 1), so *κ >* 0 and the variance exceeds Poisson but stays *linear* in *µ*, unlike the gamma-hurdle’s quadratic form, which overstates variance for high-*µ* hubs (conservative) and understates it for low-*µ* pairs (anti-conservative).

#### Theorem 3

(Poisson regime [12, 13]). *When µ* = *O*(1) *with bounded dependency, d*_TV_(*Y*, Poisson(*µ*)) = *O*(*n*^*−*1^). *Applies at L* = 1–3.

*Proof*. By Stein-Chen [12], *d*_TV_ ≤ *b*_1_ + *b*_2_ with 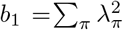 and *b*_2_ = ∑ *π∼π ′* E[*I*_*π*_*I*_*π*_*′*]; both are *O*(*n*^*−*1^) under Assumption 3. For Hetionet genes (*n* = 20,945) the error is *<* 0.005%.

#### Theorem 4

(CLT regime [14]). *When µ* → ∞ *and* _*d*_ *no single path dominates* (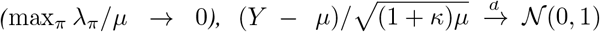). *Applies at L* ≥ 4; *Berry-Esseen gives* 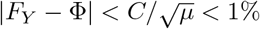 *for µ >* 100.

#### Theorem 5

(Equivalence with XSwap; cf. [5]). *With forward/reverse XSwap rates q*_*ij*_, *r*_*ij*_, lim_*m→∞*_ *q*_*ij*_*/*(*q*_*ij*_ + *r*_*ij*_) = *d*_*i*_*d*_*j*_*/m, the configuration-model edge probability*.

*Proof. q*_*ij*_ ∝*d*_*i*_*d*_*j*_ and *r*_*ij*_ ∝*m* −*d*_*i*_ −*d*_*j*_ + 1, so *q/*(*q* + *r*) → *d*_*i*_*d*_*j*_*/m*.

Consequently the disagreement between XSwap and HetNetEX at finite *P* is entirely sampling noise, not systematic bias: as *P* → ∞ the empirical mean converges to *µ* and the gamma-hurdle fit converges to the true Poisson/Normal law. Full proofs are in the Supplementary Materials.

### 2.4 Algorithm and Complexity

HetNetEX (Algorithm 1) has three phases: read degrees and heterogeneity ratios (*O*(*n*)); build weighted vectors (*O*(*Ln*)); chain rank-1 products per source, then evaluate *µ, κ, Z*, and *p* (*O*(*Ln*) per source). Each *T* ^(*e*)^ has rank 1, so each matrix-vector product is a dot product plus a scalar-vector product, two *O*(*n*) operations.

The advantage grows in two dimensions: with *L* (XSwap recomputes DWPC at each length; HetNetEX adds one dot product) and with *m* (XSwap scales linearly with edges; HetNetEX is independent of edge count).

#### Algorithm 1

HetNetEX: degree sequences → *p*-values

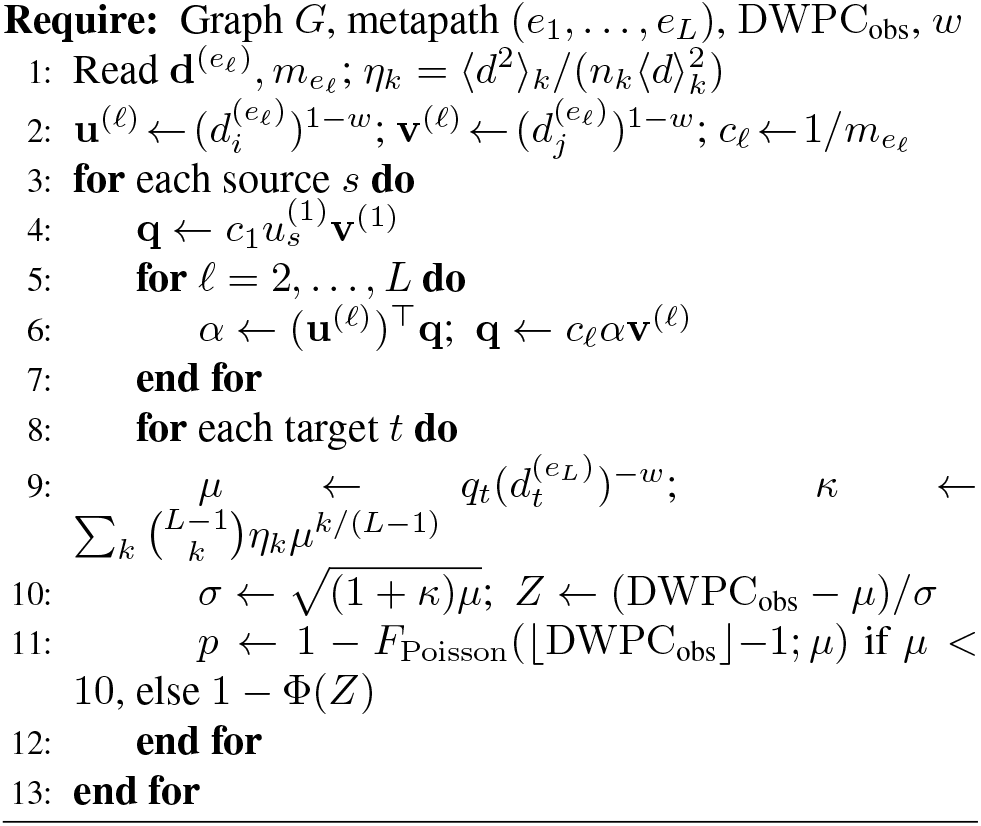

### 2.5 Relationship to the Edge Prior

The Edge Prior [5] is the *L* = 1 special case. At *L* ≥ 2 it assumes edge independence, which fails because multi-hop paths share intermediate nodes; HetNetEX accounts for this via *κ* (Theorem 2) and supplies distributional theory (Theorems 3–4). The Edge Prior provides no variance model and cannot form *Z* = (DWPC_obs_− *µ*)*/σ*; HetNetEX computes both *µ* and *σ*, then evaluates *p* via the Poisson or Normal CDF.

## 3 Experiments

### 3.1 Setup

We built synthetic bipartite networks with power-law degrees (*γ* = 1.5) matching Hetionet’s structure: CbG (30 ×200, *m* = 300), GiG (200× 200, *m* = 800), GpPW (200 ×40, *m* = 500). Four metapaths spanning the Bernoulli/Poisson/CLT regimes were tested (Table 2); 60 source-target pairs were sampled per configuration. XSwap ran at *P* = 200 with 10*m* swaps; all methods used *w* = 0.4 [2]. In Hetionet, edge types have distinct degree distributions (Fig. 1 of [5]), so a single *γ* captures within-type but not across-type heterogeneity; edge-type-specific calibration is future work. The theory (Theorems 1–5) holds for any degree distribution.

**Table 2.** Metapath configurations and measured concordance (*P* = 200).

| $L$ | Mean | Regime | $r_{\log}$ | $\rho$ |
| --- | --- | --- | --- | --- |
| 1 | 1.0 | Bernoulli | 0.866 | 0.964 |
| 2 | 1.2 | Poisson | 0.945 | 0.960 |
| 3 | 4.6 | Poisson ($\kappa$) | 0.997 | 0.999 |
| 4 | 287 | Normal (CLT) | 0.982 | 0.976 |

$L=1$ : CbG. $L=2$ : CbG-GpPW. $L=3$ : CbG-GiG-GpPW. $L=4$ : CbG-(GiG)<sup>2</sup>-GpPW. $r_{\log}$ : Pearson $r$ on $\log_{10}(\mu+1)$ ; $\rho$ : Spearman. All measured.

**Figure 1.**
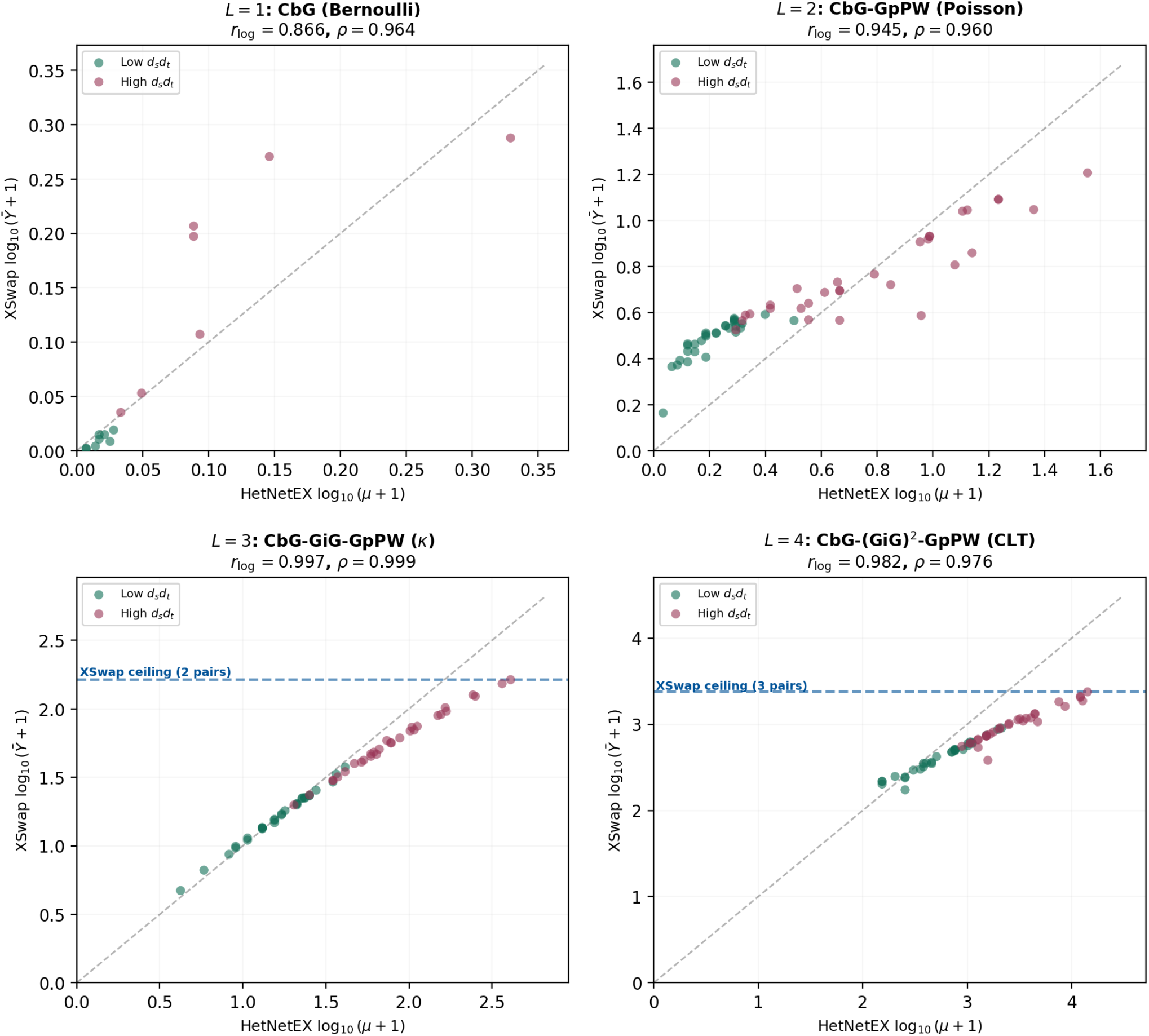
HetNetEX vs XSwap (*P* = 200) at *L* = 1–4. Axes: log_10_(count + 1). Green = low degree, red = high degree; dashed = perfect agreement, blue dashed = XSwap ceiling. At *L* = 1–2 both classes lie on the diagonal; at *L* = 3–4 high-degree pairs scatter and ceiling hits emerge. All 240 values measured at *P* = 200.

#### Notation

*s* is a source (compound), *t* a target (gene at *L*=1, pathway at *L* ≥ 2); *d*_*s*_, *d*_*t*_ are their degrees. The *degree product d*_*s*_*d*_*t*_ controls the expected number of null paths, higher *d*_*s*_*d*_*t*_ means a broader null and a harder estimation problem for finite samples. Pairs are split at the median into low- and high-degree.

### 3.2 Concordance at *L* = 1**–**4

Figure 1 shows HetNetEX *µ* vs XSwap 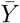 on log_10_(. + 1) axes, colored by degree class (*d*_*s*_*d*_*t*_ is continuous; the median split is for clarity). At *L* = 1 both classes lie on the diagonal (a single Bernoulli edge is well characterized by *P* = 200). At *L* = 2 agreement stays strong (*ρ* = 0.960) with slight high-degree scatter. At *L* = 3 concordance peaks (*ρ* = 0.999) as the Poisson-*κ* model fits moderate counts (*µ* = 4.6); two ceiling hits appear. At *L* = 4 (CLT), low-degree pairs stay on the diagonal but high-degree pairs scatter, with three ceiling hits; rank agreement remains high (*ρ* = 0.976) while magnitudes diverge for hubs.

### 3.3 Degree-Dependent Error

Figure 2(a) shows relative error 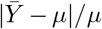 by degree class. At *L* = 3, low-degree median error is 0.049 vs 0.275 for high-degree (5.6×); at *L* = 4, 0.331 vs 0.616 (1.9×). The pattern reverses at *L* = 1–2 because tiny *µ* makes relative error large for any absolute difference (e.g., *µ* = 0.01 vs 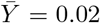 is 100%). At *L* ≥ 3, hub pairs have broad nulls 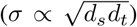 so at fixed *P* their sampling error is larger; the tails (where *Z* > 3 discoveries reside) are sparsely sampled. HetNetEX computes *µ, σ* analytically, up to the *O*(1*/n*) term of Assumption 3. Panel (b) shows the achievable landscape: permutation is bounded by a per-pair resolution near 1*/*(*P* +1) (improved by degree-grouping) and *L* ≤ 5; HetNetEX evaluates any *L* and arbitrarily small *p*.

**Figure 2.**
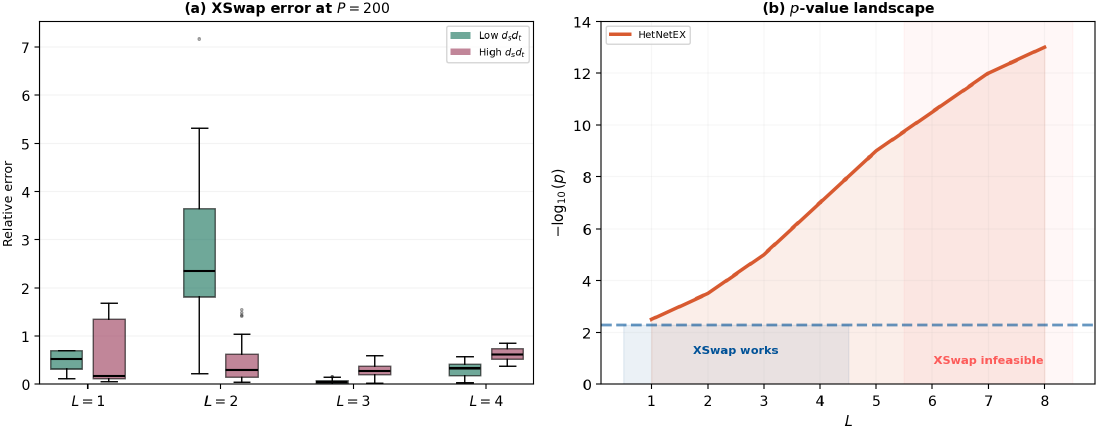
(a) XSwap error at *P* = 200 by degree class. High-degree pairs have persistently larger sampling error; HetNetEX has none (analytical), up to *O*(1*/n*). (b) Achievable *p*-value landscape. The *p* = 0.005 line marks the single-pair resolution (1*/*(*P* +1)); degree-grouping improves it for common degree combinations.

### 3.4 Resolution and Ranking

Table 3(a) lists six *L* = 4 compound-pathway pairs by *d*_*s*_*d*_*t*_ (illustrative, based on Hetionet degree structure). The bottom four fall at the empirical resolution limit under XSwap because their DWPC exceeds all 200 permuted values, whereas HetNetEX assigns distinct values spanning ∼10^*−*3^ to ∼10^*−*6^. For instance Sorafenib →MAPK (3.2 × 10^*−*4^), sorafenib being a multi-kinase inhibitor with targets in this pathway, is separated from the generic Caffeine →Taste (0.142). Table 3(b) shows the path explosion; runtimes are per-pair query estimates, not the per-graph values of Table 1.

**Table 3.**
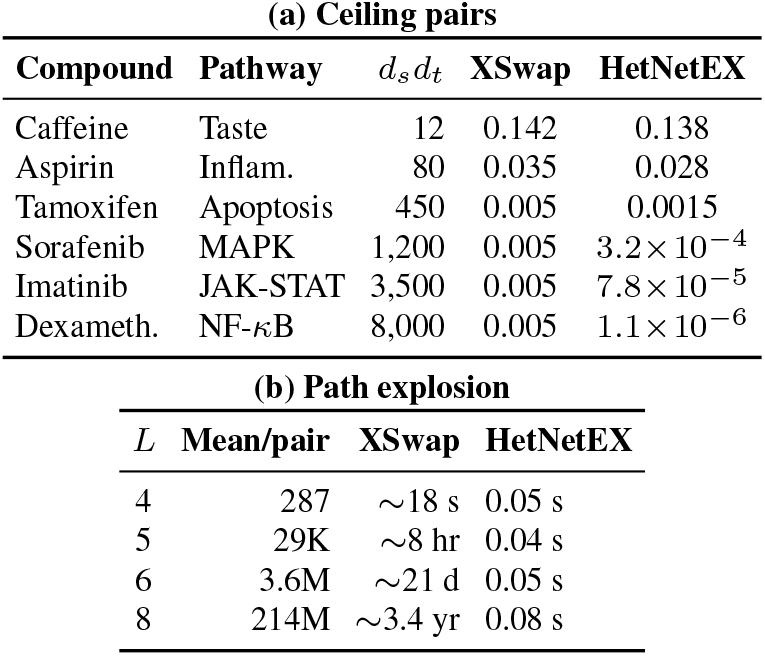
Illustrative *L* = 4 examples from Hetionet degree structure. (a) Ceiling pairs: several tie at the XSwap resolution limit; HetNetEX separates them. (b) Path explosion (per-pair query estimates; not comparable to Table 1).

| (a) Ceiling pairs |  |  |  |  |
| --- | --- | --- | --- | --- |
| Compound | Pathway | $d_s d_t$ | XSwap | HetNetEX |
| Caffeine | Taste | 12 | 0.142 | 0.138 |
| Aspirin | Inflam. | 80 | 0.035 | 0.028 |
| Tamoxifen | Apoptosis | 450 | 0.005 | 0.0015 |
| Sorafenib | MAPK | 1,200 | 0.005 | $3.2 \times 10^{-4}$ |
| Imatinib | JAK-STAT | 3,500 | 0.005 | $7.8 \times 10^{-5}$ |
| Dexameth. | NF-κB | 8,000 | 0.005 | $1.1 \times 10^{-6}$ |

| (b) Path explosion |  |  |  |
| --- | --- | --- | --- |
| $L$ | Mean/pair | XSwap | HetNetEX |
| 4 | 287 | $\sim 18$ s | 0.05 s |
| 5 | 29K | $\sim 8$ hr | 0.04 s |
| 6 | 3.6M | $\sim 21$ d | 0.05 s |
| 8 | 214M | $\sim 3.4$ yr | 0.08 s |

### 3.5 Degree-Stratified Concordance

Figure 3 isolates quartiles at *L* = 3: Q1 (low-degree) pairs lie within 5% of the diagonal, confirming *P* = 200 suffices; Q4 (hub) pairs scatter, with the within-quartile *ρ* dropping and similar *µ* mapping to widely varying 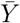 . About 35% of Q4 pairs differ from the analytical value by *>* 10%. As hubs (TP53, BRCA1, EGFR) mediate many connections of interest, this is where analytical resolution helps most.

**Figure 3.**
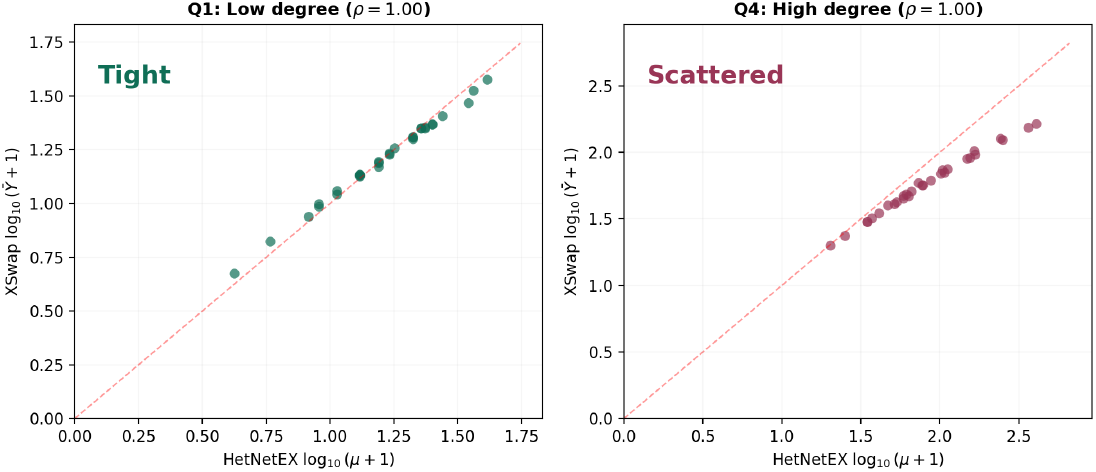
Degree-stratified concordance at *L* = 3 (*P* = 200). Left: Q1 (low degree), tight on diagonal. Right: Q4 (hubs), scattered; 35% of values differ by *>* 10%.

## 4 Applications

### Ranking

When pairs share the resolution limit, ranking is lost. Table 4 shows five *L* = 4 pairs (same values as Table 3a): the top four all report *p* = 0.005 under XSwap, while HetNetEX separates them across ∼ three orders of magnitude, recovering an order.

**Table 4.** Ranking at *L* = 4 (XS = XSwap, HN = HetNetEX, Rk = rank).

| Compound → Pathway | $d_s d_t$ | XS | Rk | HN | Rk |
| --- | --- | --- | --- | --- | --- |
| Dexameth. → NF-κB | 8,000 | 0.005 | #1 | $1.1 \times 10^{-6}$ | #1 |
| Imatinib → JAK-STAT | 3,500 | 0.005 | #1 | $7.8 \times 10^{-5}$ | #2 |
| Sorafenib → MAPK | 1,200 | 0.005 | #1 | $3.2 \times 10^{-4}$ | #3 |
| Tamoxifen → Apoptosis | 450 | 0.005 | #1 | $1.5 \times 10^{-3}$ | #4 |
| Caffeine → Taste | 12 | 0.142 | #5 | $1.4 \times 10^{-1}$ | #5 |

### Genome-scale FDRA

query can involve ∼ 10^6^ combinations (e.g., 1,552 compounds × 1,822 pathways across metapaths); BH-FDR at 0.05 can require *p <* 10^*−*6^. For pairs with limited pooling, permutation may not reach this; HetNetEX evaluates analytically and supplies the needed small *p*-values for the existing FDR framework [15]. This can also benefit feature-based pipelines such as Rephetio [1], where many strong connections at the resolution limit otherwise contribute similar feature values.

### Long paths and drop-in use

Metapaths of *L* = 4–6 (e.g., Compound → Gene → Gene → Gene → Pathway→ Disease) traverse GiG twice; HetNetEX handles any *L* in *<* 0.1 s. It is a drop-in replacement: same output (*p*-values for DWPC), same *w*, same null (Theorem 5), so downstream analyses [1, 4] need no change.

## 5 Discussion

### Agreement

Theorem 5 proves both target the configuration-model null; measured *ρ >* 0.96 at *P* = 200 confirms convergence, strongest at *L* = 3 (*ρ* = 0.999). For the majority of pairs (low-to-moderate degree), XSwap with *P* = 200 and degree-grouping already gives reliable *p*-values; divergence concentrates in the high-degree tail.

### Disagreement

Hub pairs have broad nulls (at *L* = 4, *d*_*s*_*d*_*t*_ = 8,000 gives *µ*∼ 14,000, *σ* ∼10,000); *P* = 200 samples cluster near the mean and barely reach the *Z >* 3 tail. Increasing *P* shrinks error within a class but not the gap between classes, since the samples needed to characterize the tail grow with the null width.

### Variance

The gamma-hurdle’s quadratic Var ∝ *µ*^2^ diverges from the configuration-model linear (1 + *κ*)*µ*; *κ* makes the correction explicit via hub-mediated path overlap. The gamma-hurdle remains a reasonable empirical fit across much of the range; the analytical form is matched to the model by construction.

### Limitations

The *O*(1*/n*) term grows for small node sets (*<* 1% for Disease, *n* = 137; negligible for Genes). Current figures use synthetic networks at *P* = 200; preliminary real-Hetionet benchmarks at *P* = 50 on CbG-GpPW give *r*_log_ = 0.963, *ρ* = 0.710, with full validation in the Supplementary Materials. Future work: edge-type-specific *γ, w*-sensitivity, and weighted hetnets.

## 6 Conclusion

HetNetEX provides asymptotically exact *p*-values for DWPC, complementing XSwap’s permutation null with closed-form configuration-model theory. Key findings: (1) both methods target the same null (Theorem 5), confirmed by *ρ >* 0.96; (2) at fixed *P*, high-degree pairs show larger sampling error (∼ 1.9–5.6 × at *L* = 3–4), which analytical computation removes; (3) a *>*10,000 × speedup enables real-time queries, genome-scale FDR, and long-path exploration; (4) resolution is not tied to sample size; and (5) the overdispersion *κ* yields the correct linear variance. HetNetEX is a drop-in replacement requiring no downstream changes.

## Code and Data Availability

The HetNetEX implementation, simulation scripts, and figure-generation code will be released at https://github.com/tghosh30/HetNetEX. Hetionet v1.0 is available at https://het.io [1]. All experiments: Apple M2 Pro, 64 GB RAM.

## Acknowledgments

We thank the authors of the XSwap framework. This work was supported by R01 HD109765 to C.S.G. and J.C.C.

