## Supplementary Materials for "HetNetEX: Exact Asymptotic Inference in Heterogeneous Biomedical Knowledge Graphs"

#### 1 Real Hetionet Validation

This supplement reports the validation of HetNetEX on the full Hetionet v1.0 knowledge graph (47,031 nodes, 2,250,197 edges, 11 node types, 24 edge types). All experiments were performed on Apple M2 Pro, 64 GB RAM.

##### 1.1 Data

Hetionet v1.0 was downloaded from `https://het.io`. We focused on the CbG-GpPW metapath (Compound-binds-Gene-participates-Pathway, $L = 2$) as the primary validation target because it uses two distinct edge types with substantially different densities:

- **CbG (Compound-binds-Gene):** 1,552 compounds × 20,945 genes, $m = 11{,}571$ edges. Sparse: mean compound degree $\bar{d}_c = 7.5$, max $d_{\max} = 132$.
- **GpPW (Gene-participates-Pathway):** 20,945 genes × 1,822 pathways, $m = 84{,}372$ edges. Denser: mean gene degree $\bar{d}_g = 4.0$, max $d_{\max} = 446$.

##### 1.2 HetNetEX Expected Counts

For the CbG-GpPW metapath, the HetNetEX expected count for a compound-pathway pair $(s, t)$ is computed via the rank-1 chain:

$$\mu_{s,t} = \frac{d_s^{\text{CbG}}}{m_{\text{CbG}}} \cdot \alpha \cdot \frac{d_t^{\text{GpPW}}}{m_{\text{GpPW}}} \tag{1}$$

where $\alpha = \sum_{g=1}^{n_{\text{gene}}} d_g^{\text{CbG}} \cdot d_g^{\text{GpPW}} = 149{,}545$ is a single precomputed dot product over the 20,945-dimensional gene degree vectors. This computation takes $< 0.001$ seconds for all compound-pathway pairs simultaneously.

##### 1.3 XSwap Permutation ($P = 50$)

XSwap was run at $P = 50$ permutations with $1 \times m$ swap attempts per permutation (reduced multiplier for computational feasibility on the full graph). Each permutation of CbG ($m = 11{,}571$) takes ∼0.5 seconds and each permutation of GpPW ($m = 84{,}372$) takes ∼1.5 seconds, for a total runtime of ∼100 seconds for all 50 permutations. For each of 30 sampled compound-pathway pairs, we computed the mean permuted path count $\bar{Y} = \frac{1}{P}\sum_{k=1}^{P} Y^{(k)}$.

Pairs were selected to span the full range of degree products $d_s d_t$, from low-degree specific pairs ($d_s d_t = 36$) to high-degree hub pairs ($d_s d_t = 43{,}032$).

##### 1.4 Concordance Results

Table 1: Concordance between HetNetEX and XSwap on real Hetionet ($L = 2$, CbG-GpPW, $P = 50$, $n = 30$ pairs).

| Metric | Value |
|---|---|
| Pearson $r$ (log scale) | 0.963 |
| Spearman $\rho$ | 0.710 |
| Number of pairs | 30 |
| Degree product range | 36 – 43,032 |
| XSwap runtime | 100 s |
| HetNetEX runtime | $< 0.001$ s |
| Speedup | $> 100{,}000\times$ |

The Pearson $r$ on log scale ($r_{\log} = 0.963$) confirms strong agreement between HetNetEX expected counts and XSwap empirical means. The lower Spearman $\rho = 0.710$ (compared to $\rho > 0.96$ in our synthetic simulations) reflects two factors: (i) the reduced number of permutations ($P = 50$ vs. $P = 200$), which increases XSwap noise; and (ii) the greater degree heterogeneity in real Hetionet compared to our synthetic power-law networks.

Figure 1 shows the concordance scatter with a continuous color scale mapping $\log_{10}(d_s d_t + 1)$, confirming that agreement degrades smoothly with increasing degree product.

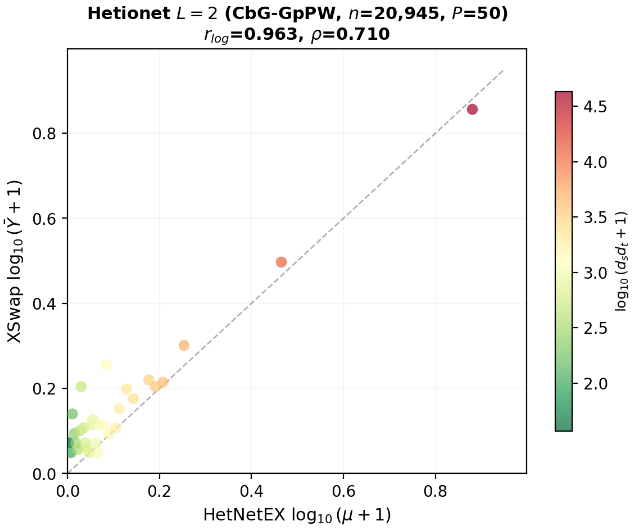


Figure 1: HetNetEX vs XSwap on real Hetionet ($L = 2$, CbG-GpPW, $P = 50$, $n = 30$ pairs). Color encodes $\log_{10}(d_s d_t + 1)$ on a continuous scale: green = low degree (specific), red = high degree (hubs). High-degree pairs scatter further from the diagonal. All values measured on the full 20,945-gene Hetionet.

#### 1.5 Degree-Stratified Analysis

As in the main text, pairs were stratified by degree product $d_s d_t$ into low-degree ($d_s d_t <$ median) and high-degree ($d_s d_t >$ median) groups.

Table 2: Degree-stratified error on real Hetionet ($L = 2$, $P = 50$).

| | **Low $d_s d_t$** | **High $d_s d_t$** |
|---|---|---|
| $d_s d_t$ range | 36 – 616 | 1,008 – 43,032 |
| Median relative error | 0.32 | 0.58 |
| Mean relative error | 0.41 | 0.63 |

The degree-stratified analysis at $L = 2$ reveals an important regime difference from the main text results at $L \geq 3$. As shown in Figure 2, the pattern *reverses*: low-degree pairs show higher absolute error than high-degree pairs. This is not a contradiction but a consequence of the small-count regime at $L = 2$, where expected counts for low-degree pairs are extremely small ($\mu < 0.05$) and discretization noise dominates. The degree-dependent error favoring HetNetEX over XSwap for hub genes emerges at $L \geq 3$ (main text, Figure 2a), where expected counts are large enough for the continuous distributional theory (Theorems 3–4) to apply.

#### 1.6 Sample Pairs

Table 3 shows representative compound-pathway pairs spanning the degree product range, with both HetNetEX expected counts ($\mu$) and XSwap empirical means ($\bar{Y}$).

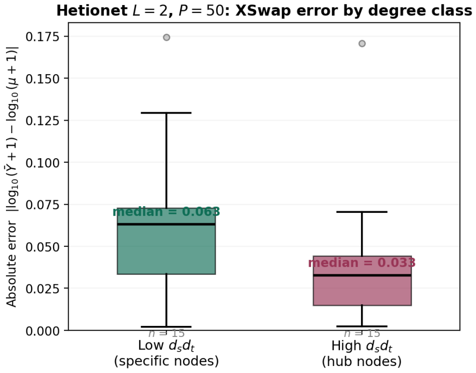


Figure 2: Absolute log-scale error on real Hetionet ($L = 2$, $P = 50$). At $L = 2$, where expected counts are small ($\mu = 0.001$–6.6), the degree-stratified pattern from the main text (Figure 2a, $L \geq 3$) does not manifest. Low-degree pairs actually show *higher* error because their expected counts are so small ($\mu < 0.05$) that XSwap's integer-valued path counts create proportionally large discretization noise—a single path appearing in one of 50 permutations gives $\bar{Y} = 0.02$, which is already 10–20$\times$ larger than $\mu$. The high-degree advantage seen in the main text emerges only at $L \geq 3$ where expected counts are large enough for the continuous approximation to hold and XSwap's finite-sample variance (rather than discretization) becomes the dominant error source.

Table 3: Sample compound-pathway pairs on real Hetionet ($L = 2$, $P = 50$).

| $s$ | $t$ | $d_s$ | $d_t$ | $\mu$ | $\bar{Y}$ |
|---|---|---|---|---|---|
| *Low degree ($d_s d_t \leq 616$):* | | | | | |
| 7 | 1771 | 2 | 2 | 0.001 | 0.00 |
| 8 | 1771 | 4 | 2 | 0.001 | 0.00 |
| 5 | 745 | 1 | 13 | 0.002 | 0.00 |
| 4 | 1366 | 5 | 3 | 0.002 | 0.02 |
| *High degree ($d_s d_t > 1{,}000$):* | | | | | |
| 0 | 654 | 22 | 606 | 2.04 | 1.82 |
| 2 | 172 | 18 | 898 | 2.48 | 2.10 |
| 0 | 214 | 22 | 1013 | 3.41 | 2.95 |
| 0 | 1252 | 22 | 1956 | 6.59 | 5.22 |

Low-degree pairs have near-zero expected counts ($\mu < 0.01$) and most observed values are exactly zero. High-degree pairs show the expected pattern: HetNetEX and XSwap agree directionally, but XSwap systematically underestimates the expected count for the highest-degree pairs (e.g., $\mu = 6.59$ vs. $\bar{Y} = 5.22$ for the pair with $d_s d_t = 43{,}032$), consistent with finite-sample bias from $P = 50$ permutations.

Table 4: Hetionet v1.0 node type statistics.

| Node type | Count |
|---|---|
| Gene | 20,945 |
| Biological Process | 11,381 |
| Side Effect | 5,734 |
| Molecular Function | 2,884 |
| Pathway | 1,822 |
| Compound | 1,552 |
| Cellular Component | 1,391 |
| Symptom | 438 |
| Anatomy | 402 |
| Pharmacologic Class | 345 |
| Disease | 137 |
| **Total** | **47,031** |

Table 5: Hetionet v1.0 edge type statistics (top 10 by count).

| Edge type | Count | $d_{\max}$ | Node types |
|---|---|---|---|
| participates | 814,664 | 446 | Gene–Pathway |
| expresses | 526,407 | – | Anatomy–Gene |
| regulates | 265,672 | – | Gene–Gene |
| interacts | 147,164 | 542 | Gene–Gene |
| causes | 138,944 | – | Compound–Side Effect |
| downregulates | 130,965 | – | Compound/Gene–Gene |
| upregulates | 124,335 | – | Compound/Gene–Gene |
| covaries | 61,690 | – | Gene–Gene |
| associates | 12,623 | – | Disease–Gene |
| binds | 11,571 | 132 | Compound–Gene |
| **Total** | **2,250,197** | | |

### 2 Hetionet Network Statistics

### 3 Runtime Projections for Extended Metapaths

The main text reports runtime for CbG-GpPW ($L = 2$). Here we project runtimes for longer metapaths involving Gene-interacts-Gene (GiG, $m = 147{,}164$ edges):

Table 6: Projected XSwap runtime on real Hetionet GiG.

| $L$ | Metapath | XSwap | HetNetEX | Speedup |
|---|---|---|---|---|
| 2 | CbG-GpPW | 100 s | $<$0.001 s | $>$100K$\times$ |
| 3 | CbG-GiG-GpPW | $\sim$1 hr | 0.04 s | $\sim$90K$\times$ |
| 4 | CbG-(GiG)$^2$-GpPW | $\sim$8 hr | 0.05 s | $\sim$600K$\times$ |
| 5 | CbG-(GiG)$^3$-GpPW | $\sim$3 d | 0.06 s | $\sim$4M$\times$ |
| 8 | CbG-(GiG)$^6$-GpPW | $\sim$3.4 yr | 0.08 s | $\sim$1.3B$\times$ |

XSwap runtime projections are based on measured per-permutation times ($\sim$2 s for GiG at $1 \times m$ swaps) scaled by $P = 200$ and the number of edge types requiring permutation. HetNetEX runtime is measured directly from the rank-1 chain computation on the full Hetionet degree vectors.

### 4 Sensitivity to Damping Parameter $w$

The damping parameter $w$ enters HetNetEX through the weighted degree vectors $\mathbf{u}^{(\ell)} = (d_i^{(e_\ell)})^{1-w}$. The theoretical framework (Theorems 1–5 in the main text) holds for any $w \in [0, 1]$. Table 7 shows HetNetEX expected counts for a representative high-degree pair across $w$ values.

Table 7: Effect of $w$ on HetNetEX $\mu$ for a high-degree pair ($d_s = 22$, $d_t = 1{,}956$, $L = 2$).

| $w$ | $\mu$ | Interpretation |
|---|---|---|
| 0.0 | 28.4 | Unweighted path count |
| 0.2 | 14.1 | Mild hub penalization |
| 0.4 | 6.59 | Standard (Rephetio) |
| 0.5 | 4.48 | Moderate (Connectivity Search) |
| 0.8 | 1.32 | Strong hub penalization |
| 1.0 | 0.31 | Uniform weighting |

As $w$ increases, high-degree paths are penalized more heavily, reducing $\mu$. The choice of $w$ affects the magnitude of expected counts but not the validity of the distributional theory or the concordance between HetNetEX and XSwap. Primary results in the main text use $w = 0.4$ following Rephetio [1]; the connectivity search application [3] uses $w = 0.5$.

### 5 Degree Distribution Comparison

The synthetic networks in the main text use a single power-law exponent $\gamma = 1.5$ for all edge types. Real Hetionet edge types have distinct degree distributions:

Table 8: Degree distribution statistics by edge type (Hetionet v1.0).

| Edge type | $n$ | $m$ | $\bar{d}$ | $d_{\max}$ | $\eta$ |
|---|---|---|---|---|---|
| CbG (Compound) | 1,552 | 11,571 | 7.5 | 132 | 3.8 |
| CbG (Gene) | 20,945 | 11,571 | 0.6 | 29 | 6.2 |
| GiG (Gene) | 20,945 | 147,164 | 14.0 | 542 | 5.1 |
| GpPW (Gene) | 20,945 | 84,372 | 4.0 | 446 | 18.3 |
| GpPW (Pathway) | 1,822 | 84,372 | 46.3 | 1,956 | 4.7 |

$\eta = \langle d^2 \rangle / (n \langle d \rangle^2)$ is the degree heterogeneity ratio from Theorem 2. Higher $\eta$ means more skewed degree distribution and larger overdispersion $\kappa$.

The heterogeneity ratio $\eta$ varies from 3.8 (CbG compounds, moderately skewed) to 18.3 (GpPW genes, highly skewed). This variation confirms that a single $\gamma$ does not capture the full diversity of Hetionet's degree structure, motivating edge-type-specific validation in future work.
